# PKIS Deep Dive Yields a Chemical Starting Point for Dark Kinases and a Cell Active BRSK2 Inhibitor

**DOI:** 10.1101/2020.06.15.153072

**Authors:** Tigist Y. Tamir, David H. Drewry, Carrow Wells, M. Ben Major, Alison D. Axtman

## Abstract

The Published Kinase Inhibitor Set (PKIS) is a publicly-available chemogenomic library distributed to more than 300 laboratories by GlaxoSmithKline (GSK) between 2011–2015 and by SGC-UNC from 2015–2017. Screening this library of well-annotated, published kinase inhibitors has yielded a plethora of data in diverse therapeutic and scientific areas, funded applications, publications, and provided impactful pre-clinical results. Based on kinome-wide screening results, we report a thorough investigation of one PKIS compound, GW296115, as an inhibitor of several members of the Illuminating the Druggable Genome (IDG) list of understudied dark kinases. Specifically, GW296115 validates as a potent lead chemical tool that inhibits six IDG kinases with IC_50_ values less than 100nM. Focused studies establish that GW296115 is cell active, and directly engages BRSK2. Further evaluation showed that GW296115 downregulates BRSK2-driven phosphorylation and downstream signaling.

**Summary Statement:** GW296115 inhibits understudied kinases, including BRSK2, with IC_50_ values less than 100nM.

## Introduction

PKIS was assembled to include 367 inhibitors designed, developed, and published by GSK. Both physical samples as well as the selectivity and potency data for PKIS were made publicly available. Compounds were chosen to provide broad coverage of the kinome, selecting diversity in chemical scaffolds and avoiding over-representation of inhibitors targeting each kinase. The composed set was distributed as a physical plate of DMSO stock solutions to all interested investigators free of charge^1^.

The National Institutes of Health (NIH) recently defined a list of proteins labeled as dark/understudied due to a lack of research and reagents to characterize their function. The IDG program was initiated to stimulate exploration of the role of these dark proteins in mediating disease initiation and propagation with the goal of providing IDG-related therapeutic avenues. One category of IDG dark proteins is kinases, where a list of 162 dark kinases was curated by the NIH and included in the IDG call for applications.

Two kinases on the IDG list, BRSK1 and BRSK2 were recently identified as inhibitors of the oxidative stress responsive transcription factor NRF2 (nuclear factor erythroid-2-related factor 2)^2^. We have reported that overexpression of active BRSK1 or BRSK2 downregulates NRF2 protein levels by suppressing protein translation. Focused experiments revealed that BRSK2 inhibits MTOR signaling while inducing phosphorylation of AMPK substrates^2^. Although overexpression experiments have demonstrated the importance of BRSK2 in cell signaling, to date, there are no reported small molecule inhibitors to characterize BRSK2 function in cells. Therefore, identifying compounds that target this kinase can yield instrumental insights into BRSK2 function.

Part of our remit for the IDG program is the creation or identification of cell active chemical tools that enable the study of dark kinases. We have taken advantage of public and internal datasets to look for potential inhibitors of IDG kinases. Through analysis of the PKIS data we identified GW296115 as a compound of interest with potent biochemical activity against a few kinases (**Fig. 1A**)^1^.

**Figure 1.**
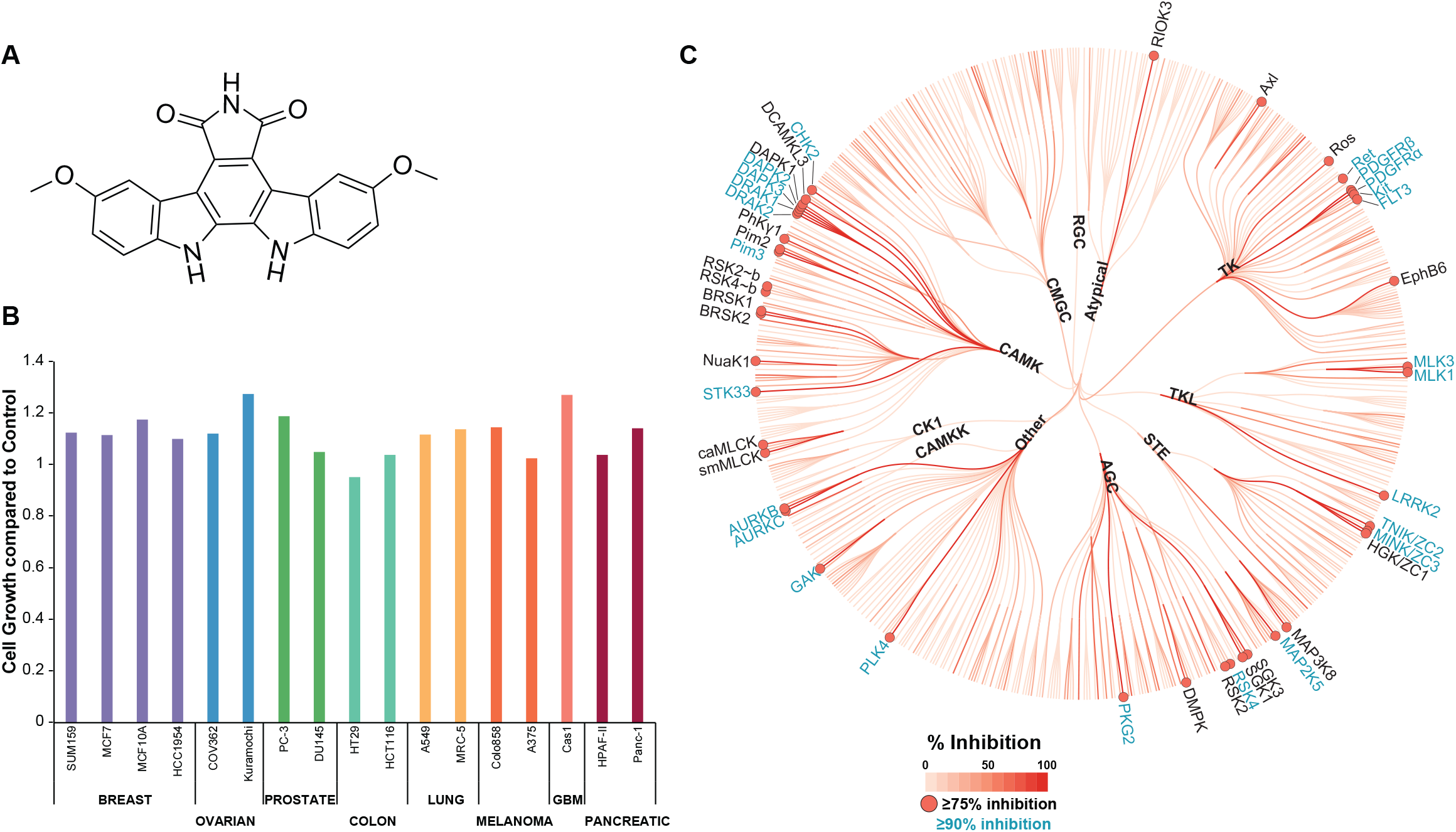
GW296115 potently inhibits kinases with minimal effect on cell growth. **A)** Structure of GW296115. **B)** GW296115 does not affect cell growth in multiple cell lines. Cell lines in the panel were treated with 1 μM GW296115 for 72h (N = 2) and analyzed using the deep dye drop method. MCF10A and MRC-5 are non-malignant cell lines. **C)** DiscoverX screen identifies 25 kinases with > 90% inhibition and 41 kinases with ≥ 75% inhibition at 1 μM of GW296115.

GW296115 is an indolocarbazole that was designed as an inhibitor of platelet-derived growth factor receptor-β (PDGFRβ). This compound, also known as 3744W, was found to inhibit the autophosphorylation of the cytoplasmic domain of PDGFR both *in vitro* (IC_50_ =1.8 ± 0.12 μM) and in insect cells (IC_50_ = 2.0 μM)^3^. Based on its activity on PDGFR, GW296115 was included in PKIS and thus screened in the Nanosyn electrophoretic mobility shift assay at two concentrations (100 nM and 1 μM). GW296115 inhibited 3 wild type human kinases >90% at 1 μM in the panel of 224 recombinant kinases (**Table S1**). These 3 kinases were also inhibited >70% at the 100 nM dose using the same assay format. Of note, PDGFRβ and PDGFRα, the originally disclosed targets of GW296115, were inhibited 85% and 84%, respectively, at 1 μM. In parallel, PKIS was profiled via thermal shift assay (TSA) against a panel of 68 kinases, of which 32 kinases were in common with the Nanosyn panel. TSA screening, which measures the increase in melting temperature (ΔT_m_) of a kinase when a small molecule inhibitor binds, identified 6 kinases with a thermal shift >7.5°C at 10 μM dose (**Table S1**)^1^. This initial screening of GW296115 highlighted 4 IDG kinases as inhibited by this scaffold: BRSK1, BRSK2, STK17B/DRAK2, and STK33.

This compound was further profiled in 17 different cancer and normal cell lines to determine whether it impacted cell growth when treated at 1 μM for 72h in duplicate. GW296115 was not found to impact the growth of any of the cell lines tested and is considered generally non-toxic to cells (**Fig. 1B**)^4^. This data aligns with the data from NCI60 panel included in the original PKIS publication, which also demonstrated that GW296115 does not impact cell growth when profiled against 60 cancer cell lines^1^. Here we further explore the target engagement characteristics of GW296115 *in vitro* and in cells, and capture its specificity for the CAMK family of kinases.

## Results

Based on its narrow profile when tested in the combined Nanosyn and TSA panels, GW296115 was included in the recently released Kinase Chemogenomic Set (KCGS). Since its inclusion in KCGS, another broader round of screening was carried out at DiscoverX. GW296115 was profiled against 403 wild type human kinases at 1 μM using the DiscoverX *scan*MAX screening platform, which employs an active site-directed competition binding assay to quantitatively measure interactions between test compounds and kinases^4,5^. GW296115 inhibited 25 kinases >90% at 1 μM, resulting in a selectivity index (S_10_) of 0.062 at 1 μM (**Table S2**, **Fig. 1C**) ^6^. However, GW296115 did not meet our criteria (S_10_(1 μM) < 0.04) for follow-up K_D_ measurement on kinases inhibited >80% at 1 μM^4^.

Since GW296115 did not meet our criteria for K_D_ follow-up at DiscoverX, we opted to collect full dose-response curves for all wild type kinases inhibited ≥75% in the DiscoverX panel that were offered by Eurofins. Eurofins has kinase enzymatic radiometric assays for 35 of the kinases on this list (**Table S2**) and a kinase enzymatic LANCE assay for one additional kinase (MAP2K5/MEK5). All assays were carried out at the K_m_ of ATP and in dose-response for the radiometric assays (9-point curve in duplicate), while the LANCE assay was carried out at a single concentration (10 μM) in duplicate (**Table S2**, **Fig. 2A**). Out of the 35 kinases evaluated, 6 IDG kinases demonstrated IC_50_ < 100nM in the Eurofins enzymatic assays (**Fig. 2B – 2G**).

**Figure 2.**
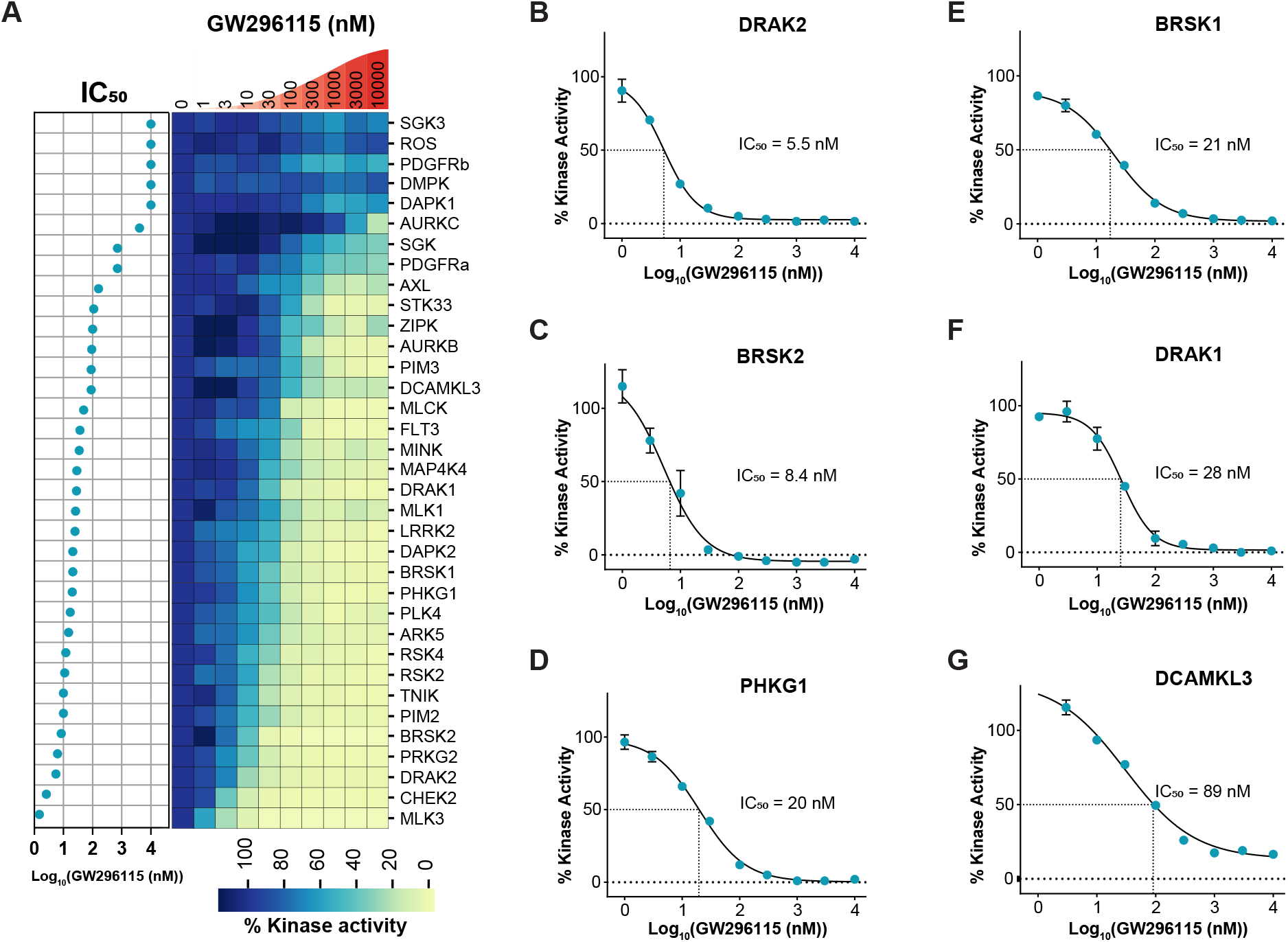
Dose-response kinase enzymatic activity assays on 35 kinases treated with GW296115 reveal its biochemical potency. **A)** Heatmap of % kinase activity with 9-point dose response to GW296115, plotted in descending order of IC_50_ values from Eurofins screen (N = 2). **B-G)** Dose response curves for IDG kinases with IC_50_ < 100 (N = 2): DRAK2 (IC_50_ = 5.5 nM), BRSK2 (IC_50_ = 8.4 nM), PHKG1 (IC_50_ = 20 nM), BRSK1 (IC_50_ = 21 nM), DRAK1 (IC_50_ = 28 nM), and DCAMKL3 (IC_50_ = 89 nM), respectively.

Our interest in illuminating BRSK2 biology motivated further investigation of the cellular target engagement of BRSK2 by GW296115 using our NanoBRET assay. BRSK2 kinase was fused to 19-kDa luciferase (NLuc) at its N-terminus (NLuc–BRSK2) which was transiently expressed in HEK293 cells, and then incubated with a cell-permeable fluorescent energy transfer probe (tracer)^7^. Using increasing concentrations of GW296115, dose-dependent displacement of tracer was observed, and provided us with an in-cell IC_50_ value. Cellular target engagement of BRSK2 in live cells was observed with an IC_50_ = 107 ± 28 nM, confirming GW296115 could be considered a potent, cell active compound (**Fig. 3A**).

**Figure 3.**
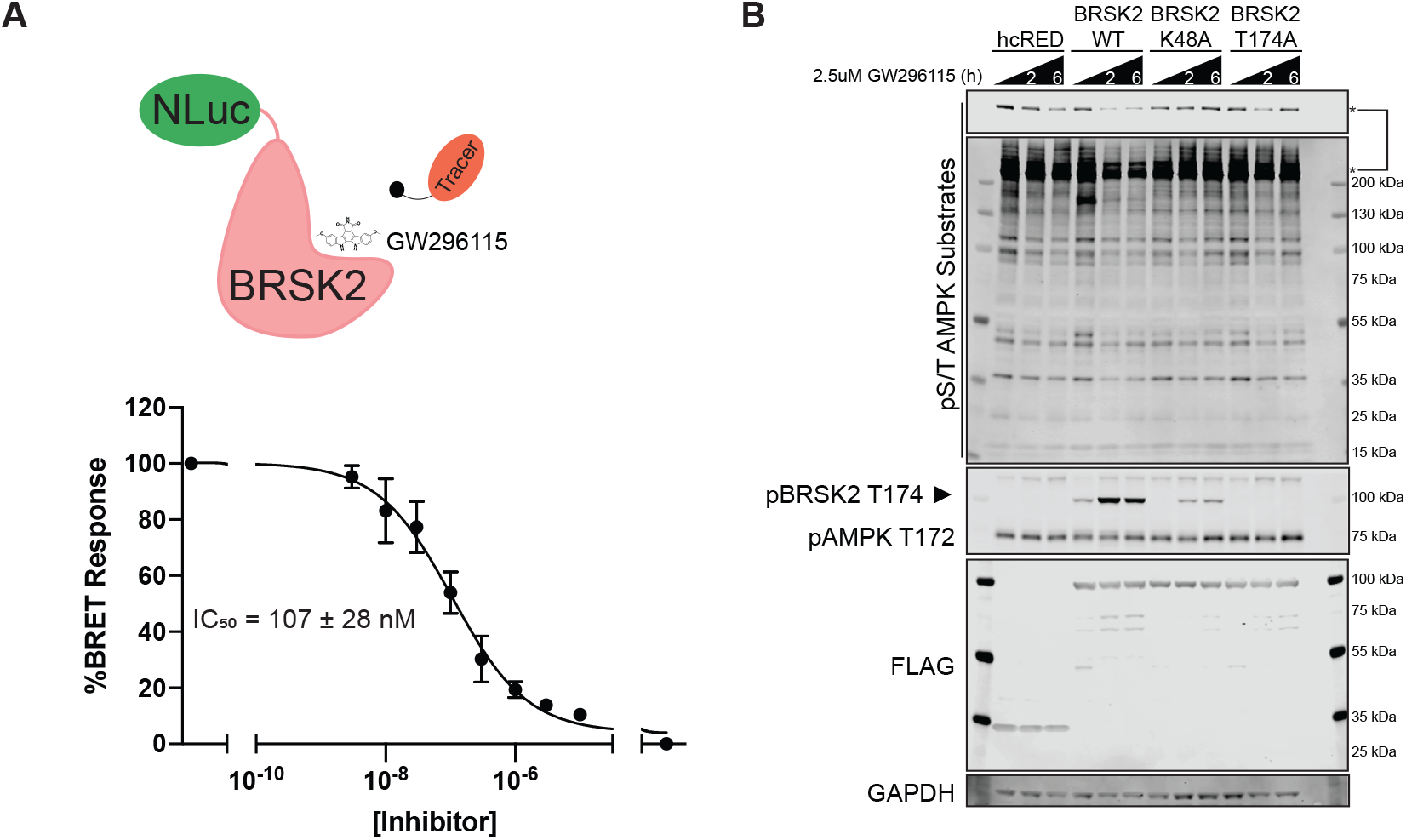
GW296115 inhibits BRSK2 in cell-based assays. **A)** GW296115 engages BRSK2 in cells with an IC_50_ = 107 ± 28 nM. HEK293 cells were transiently transfected with NLuc-BRSK2 and GW296115 target engagement was calculated by measuring the dose-dependent displacement of the tracer (N = 3). **B)** GW296115 inhibits kinase active BRSK2 and downregulates AMPK substrate phosphorylation. HEK293T cells transiently expressing hcRED, wild type (WT), or kinase dead (K48A & T174) were treated with 2.5 μM of GW296115 for 2 or 6 hours. Unformatted images of blots are included in **Fig. S2**.

With confirmed cellular activity, we next focused on characterizing the pathways mediated by BRSK2 utilizing GW296115. BRSK2 has a similar kinase domain to AMPK and is activated by LKB1-mediated phosphorylation in the active site at T174^8–10^. As previously reported, BRSK2 induces phosphorylation of AMPK substrates^2^. Therefore, we asked whether this BRSK2-induced phosphorylation of AMPK substrates is ablated by GW296115. First, we evaluated the effect of GW296115 on AMPK substrate phosphorylation in HEK293T cells overexpressing wild-type or kinase-dead BRSK2 (**Fig. 3B**). Specifically, we overexpressed two different kinase-dead BRSK2 constructs, K48A and T174A, and treated cells with 2.5μM of GW296115 for 2 or 6 hours. Compared to cells expressing hcRED control, wild-type BRSK2 overexpression induced AMPK substrate phosphorylation, as measured by phospho-S/T AMPK substrate antibody. This change was not observed when overexpressing kinase-dead variants. BRSK2-induced AMPK substrate phosphorylation was ablated with GW269115 treatment at both 2- and 6-hour time-points.

Moreover, we used an antibody against pAMPK T172, a known LKB1 target site, and found that phosphorylation at T172 was not altered. However, using the same antibody we were able to detect phosphorylation of BRSK2 at T174, which showed hyper-phosphorylation in response to GW269115 in all samples except BRSK2 T174A expressing cells.

Next we evaluated whether specific *bona fide* AMPK substrates were phosphorylated by BRSK2. Based on the total pS/T AMPK substrate blots, we decided to check known substrates that match the size of the most robust changes due to BRSK2 overexpression. UNC51-like kinase 1 (ULK1) is a 120kDa kinase that is member of the autophagy initiation complex, and is phosphorylated by AMPK at multiple residues, S317 and S555, among others^11–13^. Therefore, we overexpressed wild type BRSK2 in HEK293T cells and measured changes in pULK1 S317 and S555 following treatment with increasing doses of GW296115 for 2 hours. BRSK2 overexpression increased phosphorylation of ULK1 at S317, but not S555, which was decreased dose-dependently by GW296115 (**Fig. 4A, 4B**). Total AMPK levels remained unchanged and western blots using pAMPK T172 showed increased levels of pBRSK2 T174, but the levels of AMPK phosphorylation were not discernable due to masking by BRSK2 overexpression (**Fig. 4A**). However, in samples expressing control hcRED, treatment with GW296115 increased pAMPK T172. We also asked if activating phosphorylation of ULK1 leads to increased phosphorylation of downstream components of the autophagy complex. Therefore we evaluated phosphorylation of S351 on P62 (SQSTM1), which is a stress induced autophagy receptor for ubiquitylated cargo^14^. Due to its central role as a signaling hub, P62 accumulation and phosphorylation serves as a sensor for starvation, oxidative stress, and selective autophagy^14–16^. Following BRSK2 overexpression, we observed increased pP62 S351, which is dose dependently downregulated by GW296115 (**Fig. 4A, 4C**). The total P62 expression level was not significantly altered in response to GW296115. Overall, these data show that BRSK2 induced AMPK substrate phosphorylation including ULK1 and the downstream autophagy effector P62. Moreover, these phosphorylation events were ablated by GW296115 in a dose dependent manner (**Fig. 4B, 4C**).

**Figure 4.**
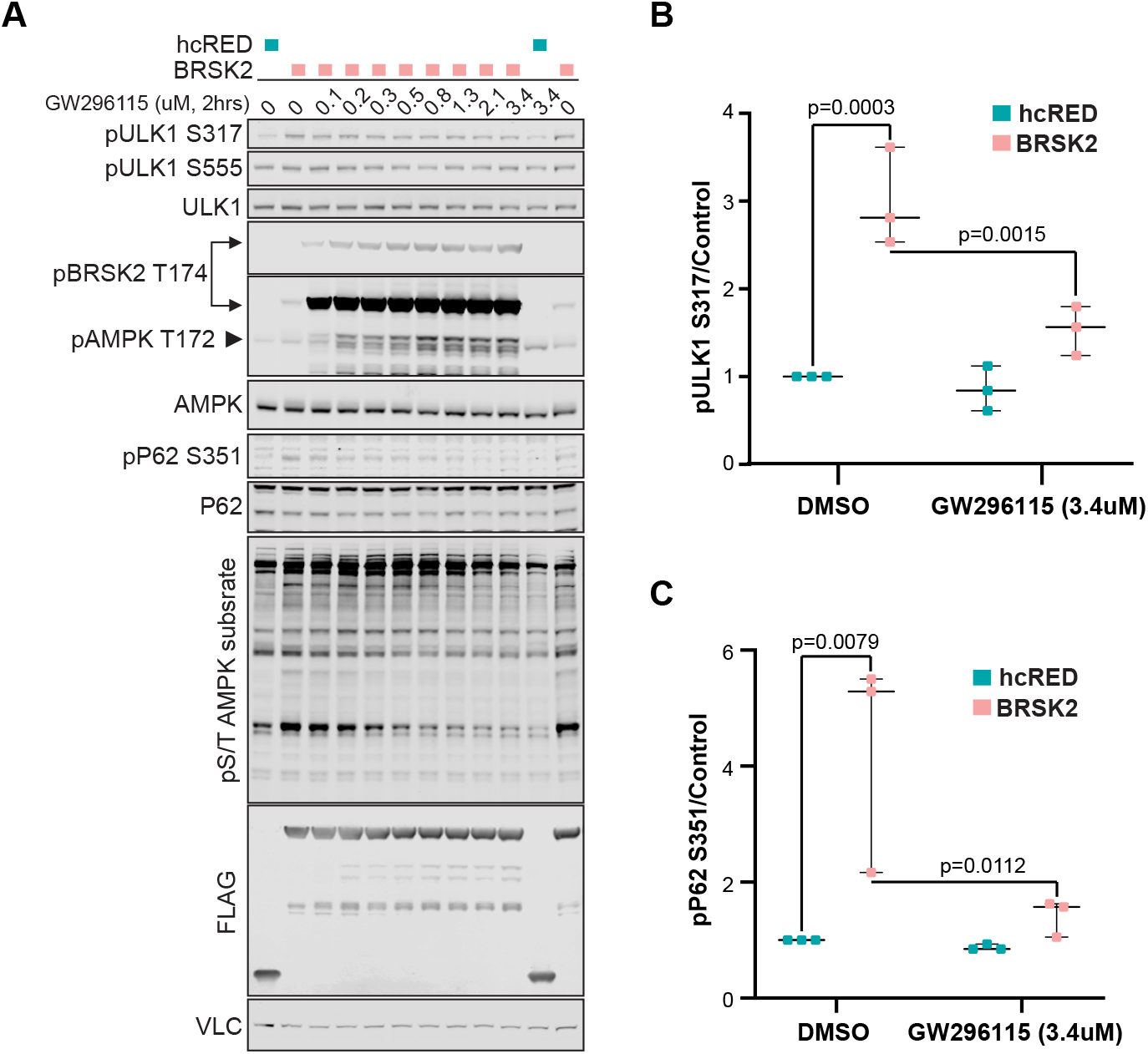
GW296115 dose-dependently inhibits BRSK2-induced phosphorylation. **A)** BRSK2 overexpression induced pULK1 S317, pP62 S351, and pS/T AMPK substrates are decreased dose-dependently by GW296115. HEK293T cells were transiently transfected with hcRED or BRSK2 for 24 hours before GW296115 treatment for 2 hours. Unformatted images of blots are included in **Fig. S2**. **B, C)** Western blot quantitation for pULK1 s317 and pP62 S351 treated with DMSO or GW296115 at 3.4 μM shows statistically significant changes (N = 3).

Finally, to inform future analog design, we selected a panel of structurally related indolocarbazoles and bisindolylmaleimides to profile in the BRSK2 NanoBRET assay (**Fig. 5A**). Kinome-wide selectivity as well as biochemical potency on BRSK2 and related CAMK family kinases has been published for all except Arcyriflavin A and K-252c^17,18^. Where available, the S_10_ selectivity score corresponding to the percentage of kinases inhibited >90% at the concentration shown is included for each compound versus GW296115 (**Fig. 5B)**. The average of two replicates shown as percent remaining kinase activity in the presence of 0.5 μM compound relative to solvent control is shown for each CAMK kinase (**Fig. 5D)**^17^. Although the assay formats are different, including the concentration of compound added and the protein constructs employed, our profiling of GW296115 is included for comparison purposes (% control at 1 μM) **(Fig. 5D)**. Given its weak biochemical inhibition of BRSK2, bisindolylmaleimide IV was excluded from testing in the NanoBRET assay. Bisindolylmaleimide I and Gö 6983, like bisindolylmaleimide IV, are highly UV active compounds and their UV absorbance interfered with the assay readout. With the exception of SB 218078, the rank order in terms of biochemical potency and cellular target engagement of BRSK2 by the compounds tracked (**Fig. 5B, S1**). For some, biochemical and cellular potency matched well, while for others there was a modest loss in potency when moving into cells. We were able to confirm that structurally related compounds, Arcyriflavin A and K-252c, for which BRSK2 biochemical inhibition data was not found, engage BRSK2 in cells. Overall, the BRSK2 NanoBRET data supports that indolocarbazoles and bisindolylmaleimides enter cells and potently engage with BRSK2. Overall these data support that GW296115 is the most selective of the potent and cell-active indolocarbazole and bisindolylmaleimide BRSK2 inhibitors profiled.

**Figure 5.**
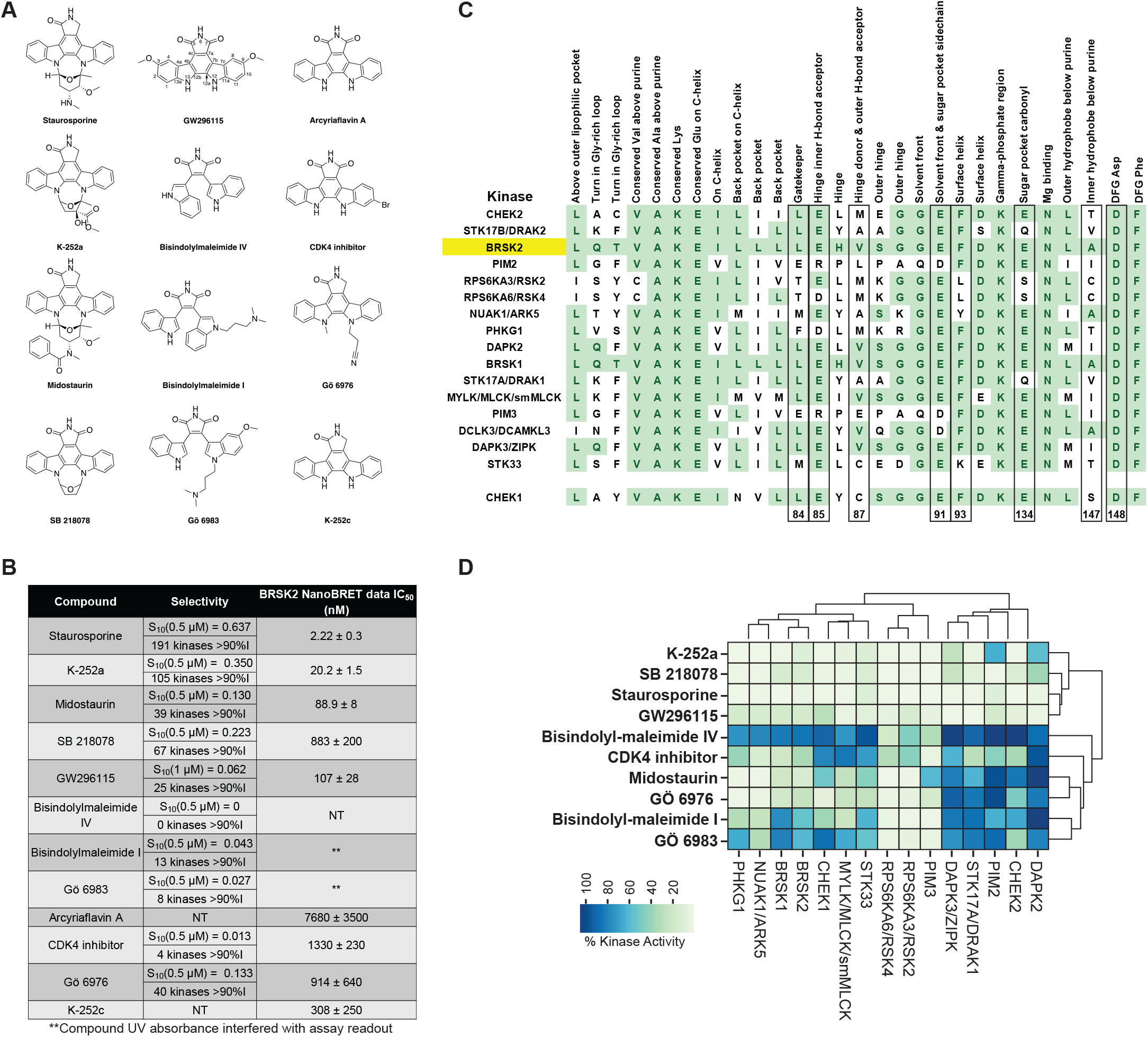
Structural- and activity-based comparisons of GW296115 to related compounds. **A)** Structures of staurosporine, K-252a, midostaurin, SB 218078, bisindolylmaleimide IV, bisindolylmaleimide I, Gö 6983, Arcyriaflavin A, CDK4 inhibitor, Gö 6976, and K-252c versus GW296115. **B)** Selectivity of compounds shown in panel A when profiled against 300 (staurosporine, K-252a, midostaurin, SB 218078, bisindolylmaleimide IV, bisindolylmaleimide I, Gö 6983, CDK4 inhibitor, and Gö 6976) or 403 (GW296115) wild type human kinases and NanoBRET data for most inhibitors in panel A (N = 3). **C)** Alignment of key residues corresponding to CAMK kinases potently inhibited by GW296115 and CHEK1. Residues colored green demonstrate homology shared with BRSK2 and residue numbers listed below correspond with those discussed with respect to SB 218078 binding to CHEK1 (also boxed). **D)** Inhibition data corresponding with those CAMK kinases (and CHEK1) included in profiling of inhibitors listed in panel A.

We can examine key residues in the respective binding pockets and use co-crystal structures to begin to rationalize why certain kinases respond differently to these structurally related compounds. Looking across residues important for binding, we see a great deal of conservation within the CAMK kinases inhibited by GW296115 (**Fig. 5C**). All kinases are compared to BRSK2 and conserved residues are highlighted in green. While CHEK1 was inhibited 70% at 1 μM by GW296115 (**Fig. 5D**), it is included because the structure of related compound, SB 218078, was solved and published with the CHEK1 kinase domain (PDB: 1NVS) and thus its homology with other CAMK family members is relevant^19^. SB 218078, like staurosporine, is a potent CHEK1 inhibitor. While both SB 218078 and staurosporine were found to bind in the same pocket, the side-chain conformations of residues E^91^, F^93^, E^134^, S^147^, and D^148^ varied between the two structures and SB 218078 was found to make no direct contacts with E^91^ and E^134^ (**Fig. 5C**). This finding supports that a ring system connecting nitrogens N12 and N13, like that found in staurosporine, is able to make favorable interactions with the binding pocket in many kinases, increasing binding affinity and inhibition potential. Furthermore, E^91^ and E^134^ have been characterized as defining the sugar pocket when ATP binds (**Fig. 5C**), supporting that the N12-N13 ring system on staurosporine fits into the same pocket as the sugar on ATP. The maleimide of SB 218078 makes hydrogen-bonding interactions with the backbones of E^85^ and C^87^, two residues that have been characterized as making key hydrogen bonds with the hinge-binding part of the molecule (**Fig. 5C**). Compared with staurosporine, SB 218078 was found to shift slightly outward from the pocket to avoid close contact between the 7-keto oxygen (C7 carbonyl) and the side chain of L^84^ (the gatekeeper residue). Based on the binding orientation of SB 218078 to CHEK1, with the N12-N13 ring system oriented toward S^147^ and D^148^, it is likely that these residues are too far away to make key interactions with GW296115 upon binding^19^. Loss of these interactions when GW296115 binds to CHEK1 is proposed to result in its loss of affinity for CHEK1 versus SB 218078 and staurosporine (**Fig. 5D**).

Through analysis of the binding of SB 218078 to CHEK1, we have identified several putative interactions that GW296115 makes with CAMK kinases, which share many highly conserved residues. IDG kinases that are potently inhibited by GW296115 (BRSK1, BRSK2, DRAK1, and DRAK2) share common features of their binding pocket, which include a polar/charged residue (Q or K) within the glycine-rich loop and aliphatic residues at positions 87 and 147 (V or A). This pattern is unique to these kinases amongst the larger CAMK kinases inhibited by GW296115. The DAPKs, which are the most homologous kinases to DRAK1 and DRAK2, and NUAK1, which is among the most homologous kinases to BRSK1 and BRSK2, show overlap in several, but not all of these residues (**Fig. 5C**). Based on the co-crystal structures of SB 218078 and staurosporine bound to CHEK1, we know that positions 87 and 147 make key interactions when this class of molecules are bound^19^. When comparing the sequence of CHEK1, which is less potently inhibited by GW296115, with those CAMK family members more potently inhibited, it appears that the back pocket residues in addition to residue 147 are most different. CHEK1 has an arginine in the back pocket on the C-helix and a serine at residue 147. An overlapping residue at these two positions is not found when surveying those CAMK kinases potently inhibited by GW296115 (**Fig. 5C**). This finding once again points to the inner hydrophobic residue in the pocket that binds the purine core of ATP (147) as making key interactions and potentially dictating binding affinity. Further, the co-crystal structures support that the side-chain conformation of residue 147 varies between structures^19^. Through taking advantage of differences in key residues within the binding pocket, we can design molecules that are more selective for subfamilies or even specific kinases of interest.

## Discussion

The publicly available PKIS library provides a platform to identify tool compounds for a number of kinases, including the 162 categorized as understudied in the NIH IDG effort. Developing better reagents to evaluate these kinases will help define poorly characterized signaling networks. Here, we show that GW296115 is a potent inhibitor of many kinases. GW296115 was included in the recently released KCGS based on profiling against 260 human kinases. The broader round of screening carried out at DiscoverX included 403 wild type human kinases. More comprehensive profiling revealed GW296115 to be much less selective than the original data suggested. The difference in selectivity observed between the Nanosyn and DiscoverX panels highlights the value of utilizing multiple orthogonal assay formats to assess compounds and problems that can arise if too much emphasis is placed on one readout.

To ascertain selectivity of GW296115 and validate our DiscoverX screening results using an orthogonal assay format, we chose to execute Eurofins enzymatic kinase assays in dose-response at the K_m_ of ATP. It is worth noting, however, that inhibition of several kinases can elicit polypharmacology and has been exploited to develop kinase drugs for cancer. Bosutinib, ponatinib, sunitinib, cabozantinib, and dasatinib are some examples of FDA-approved kinase drugs that are less selective than GW296115, with S_10_(1 μM) scores of >0.15 when tested against 305–311 kinases at 1 μM. These 5 drugs approved for clinical use were profiled using a combination of mobility shift assays and ELISA technology, keeping the ATP concentration within 2-fold of the K_m_ of ATP for every individual kinase^20^. While the methods used to profile GW296115 and these drugs vary, as do the size of kinase panels, it is clear that kinome-wide selectivity of kinase-targeting compounds is not the only determinant in assessing their utility.

We found that 25 kinases that were profiled at Eurofins exhibited an IC_50_ < 150 nM, where 8 kinases had IC_50_ ≤ 10nM (**Fig. 6**)^6,21^. Importantly, 6 kinases with an IC_50_ < 100 nM are IDG dark kinases (RIOK3, also dark, was not tested at Eurofins). The 4 IDG kinases identified via Nanosyn and TSA screening (BRSK1, BRSK2, STK17B/DRAK2, and STK33) were once again captured in the DiscoverX profiling and confirmed as potently inhibited by GW296115 at Eurofins. In fact, 8 of 9 kinases from **Table S1** were also inhibited ≥75% at DiscoverX. PRKAA2 is the outlier as treatment with GW296115 at 1 μM resulted in 20% inhibition in the DiscoverX *scan*MAX panel. Comparison of **Tables S1** and **S2** once again demonstrates the power of using orthogonal assay formats for data generation to identify overlapping data and discern possible misleading hits.

**Figure 6.**
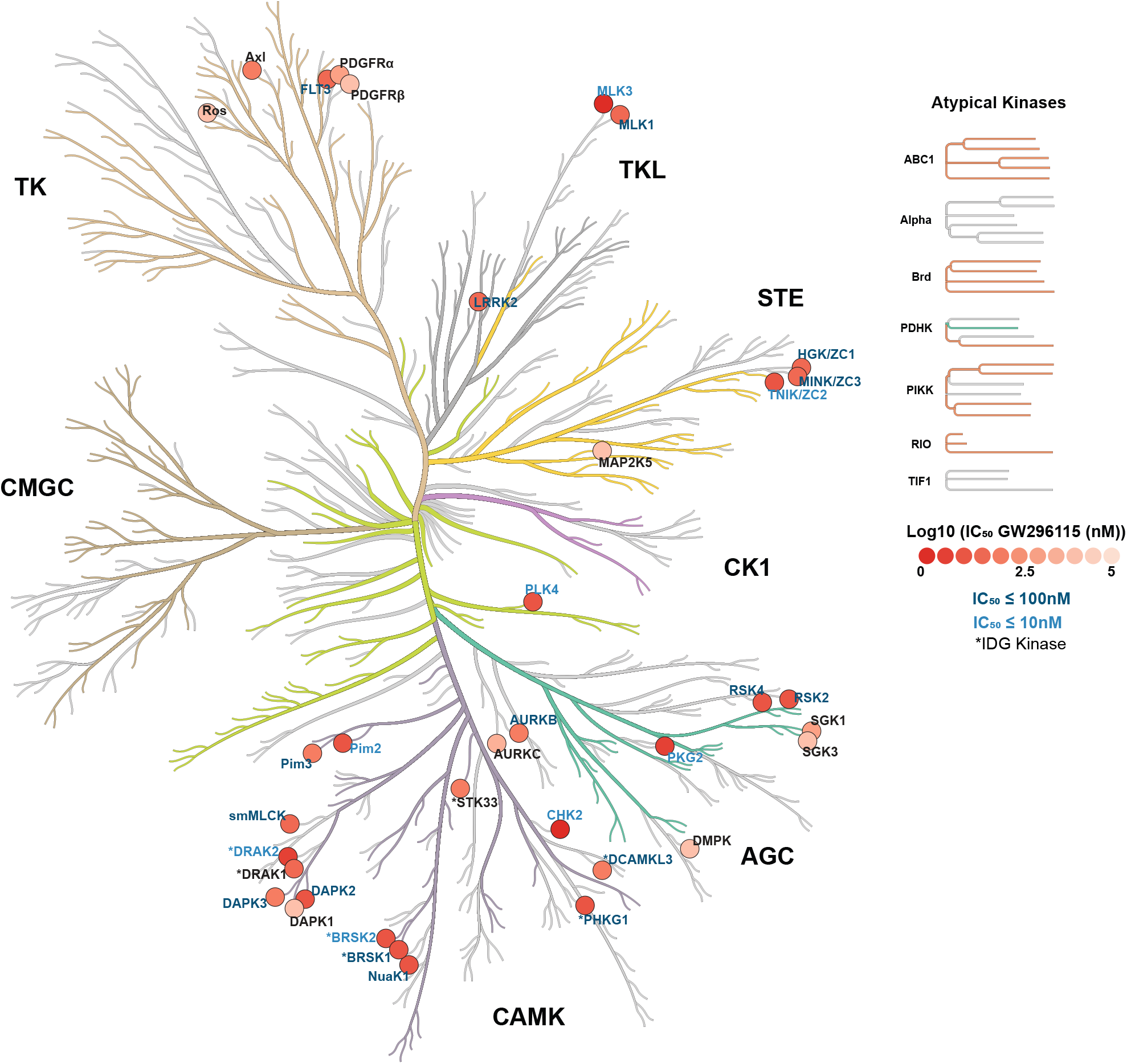
Enzymatic assay profile of GW296115. CORAL kinome tree representing kinases inhibited by GW296115 labeled according to IC_50_, and IDG status (*).

Building on the biochemical data generated for GW296115, we carried out studies to determine whether this compound was active in cells. Given its confirmed potency against BRSK1/2 in the Nanosyn, DiscoverX, and Eurofins panels and our interest in these dark kinases, we first looked to evaluate GW296115 for cellular target engagement and pathways regulated by BRSK2. Utilizing BRSK2 NanoBRET assay we confirmed that GW296115 permeates live cells and exhibits potent engagement of BRSK2 with an IC_50_ = 107 ± 28 nM.

With confirmed cell activity, we moved to cell-based experiments to probe how this compound impacts BRSK2-mediated pathways. GW296115 inhibited phosphorylation of AMPK substrates in BRSK2-overexpressing HEK293T cells. Moreover, this effect was observed after just 2 hours of treatment. This finding confirms GW296115 as a cell-active tool that interrupts a phosphorylation events driven by BRSK2. While GW296115 did not inhibit phosphorylation at the LKB1 target site (T172) on AMPK, BRSK2 was hyper-phosphorylated on T174 in response to GW296115. BRSK2 T174 hyper-phosphorylation was more notable when overexpressing wild-type BRSK2, but was also observed when overexpressing K48A BRSK2. This later observation supports that GW296115 may result in activation of a compensatory mechanism that induces hyper-phosphorylation when BRSK2 is inhibited, possibly through activation of feedback loops. Further probing of well-defined AMPK substrate ULK1 and downstream effector P62 demonstrated that GW296115 inhibited BRSK2-induced phosphorylation in HEK293T cells in a dose-dependent manner. These studies establish that GW296115 can be used to interrupt BRSK2-mediated pathways in cells.

Stemming from interest surrounding the natural product staurosporine, a plethora of chemistry has centered on the bisindolylmaleimide scaffold. Staurosporine (**Fig. 5A**) is one of the most potent and non-selective kinase inhibitors known. Analogs have been designed to reduce the structural complexity of the natural product while, at the same time, decrease the number of kinases it potently inhibits. Exemplars in **Fig. 5A**, like GW296115, bear a maleimide to adjoin the two indole ring systems. All inhibitors, with the exception of Arcyriaflavin A, K-252c, and GW296115, were profiled against a panel of 300 recombinant protein kinases at 0.5 μM. These were 9 of the 178 commercially available kinase inhibitors included in this study by Anastassiadis and co-workers. Staurosporine, K-252a, and SB 218078 were found to be the top 3 most promiscuous of all compounds tested, respectively (**Fig. 5B**)^17^. Through examination of structure versus selectivity, we can clearly see that the bisindolylmaleimide scaffold is tunable. Removal of the carbon-carbon bond between C12a and C12b results in great losses in potency across the kinome, suggesting that free rotation of the two indole ring systems is not well tolerated by many kinases and may prevent bisindolylmaleimides IV and I as well as Gö 6983 from making key interactions with the ATP binding site. Addition of the C-7 carbonyl is also tolerated by some kinases but not others. The ring system that connects nitrogens N12 and N13 seems to make favorable interactions with kinase binding sites such that more kinases are potently inhibited by inhibitors that have it (staurosporine, K-252a, and SB 218078) versus those that do not (GW296115 and CDK4 inhibitor, for example). The pocket that binds this N12-N13 ring in many kinases, however, does not tolerate too much bulk, so midostaurin with its N-benzoyl group likely results in unfavorable steric clash. N-alkylation of the indolyl nitrogens is done to attempt to access binding interactions with the part of the pocket that accommodates this N12-N13 ring and the sugar on ATP. Finally, substitution at C3, C9, and/or C10 also impacts the number of kinases potently inhibited by bisindolylmaleimides/indolocarbazoles. This finding is likely due to the steric clash introduced by these groups with binding pocket residues, which is tolerated by some kinases but prevents binding to others.

Our screening data generated around GW296115 demonstrates that the CAMK family of kinases is particularly sensitive to this compound and many members are potently inhibited by it. The majority of kinases that demonstrated an IC_50_ <115 nM in the Eurofins radiometric and LANCE assays are from the CAMK family (**Table S2, Fig. 6**). These CAMK family kinases are listed from most to least potently inhibited by GW296115 in **Fig. 5C**. When Anastassiadis and co-workers profiled their 178 commercial inhibitors, several of these CAMK family members were included in the 300-kinase panel. The same trend was observed within the CAMK family as when overall selectivity was calculated (**Fig. 5B**): the most CAMK kinases were inhibited by staurosporine and the least by bisindolylmaleimide IV (**Fig. 5D**)^17^. As discussed above, certain CAMK kinases are more sensitive to structural changes than others. RSK2 and RSK4, for example, are potently inhibited by all but bisindolylmaleimide IV and CDK4 inhibitor. In contrast, the DAPK2 ATP binding site is very sensitive to structural changes, which is evident when considering inhibition data for staurosporine versus closely related K-252a.

All structurally related indolocarbazoles and bisindolylmaleimides with the exception of bisindolylmaleimide IV were tested in the BRSK2 NanoBRET assay to evaluate their cellular target engagement of BRSK2 in live cells (**Fig. 5A**). While bisindolylmaleimide IV was excluded due to its weak biochemical potency on BRSK2, it is also highly UV active. Bisindolylmaleimide I and Gö 6983 are two additional highly UV active compounds in the series. Testing all 3 compounds using the NanoBRET format was confounded by the high UV absorbance, which interfered with the assay at high concentrations (**Fig. S1**). When we compare the NanoBRET data (**Fig. 5B**) with the biochemical data (**Fig. 5D**), we observe the same potency trend for compounds included in both panels with the exception of SB 218078. For staurosporine, K-252a, and midostaurin, potent inhibition in an enzymatic assay translated to low nanomolar engagement of BRSK2 in the NanoBRET assay (IC_50_ <100 nM). While these compounds were more active than GW296115 in the BRSK2 NanoBRET assay, they are much less selective than GW296115 when profiled broadly (**Fig. 5B**). Interestingly, K-252c, for which reported biochemical BRSK2 data was not published, was found to engage BRSK2 with an IC_50_ = 308 ± 250 nM. Arcyriflavin A, which was also devoid of BRSK2 biochemical data in the literature and only differs from K-252c at the C-7 position, was found to be much less active (IC_50_ = 7680 ± 3500). Like the remainder of the compounds tested in the BRSK2 NanoBRET assay that seem like outliers and have IC_50_ values in the 600–1600 nM range (SB 218078, Gö 6976, CDK4 inhibitor), Arcyriflavin A suffers from poor solubility in DMSO, poor cell penetrance, and/or UV interference in the NanoBRET assay. A combination of these factors explain the poor translation of biochemical activity to cell-based potency. The BRSK2 NanoBRET data indicates that GW296115 and related structures can enter cells and bind to BRSK2, but that compound solubility and/or cell penetrance is a challenge that must be overcome with design.

These structure-based observations and corresponding inhibition data around GW296115 and related bisindolylmaleimides/indolocarbazole support the idea that these scaffolds can be tuned to improve kinome-wide selectivity. Kinases are clearly sensitive to specific modifications within these scaffolds. The dataset shown in **Fig. 5** supports GW296115 as the most potent and selective, cell-active inhibitor of BRSK2 within the structural class.

## Conclusions

GW296115 represents a potent, cell active chemical starting point from which we can design inhibitors. The promiscuity of this compound is clearly a feature we can exploit. Among the kinases that were potently inhibited in corresponding enzymatic assays are BRSK1 (IC_50_ = 21 nM), BRSK2 (IC_50_ = 8.4 nM) and near neighbor (based on homology) NUAK1 (IC_50_ = 15 nM). The biochemical potency of this compound translates in cells, as it demonstrated activity in both the NanoBRET and phosphorylation assays. While GW296115 cannot be considered selective, it is the best available chemical tool to study BRSK1/2 biology.

## Materials and methods

### Kinome screening

The *scan*MAX assays were performed at Eurofins DiscoverX Corporation as previously described^5^.

### *In vitro* kinase radiometric and LANCE assays

Eurofins kinase enzymatic radiometric assays were carried out at the K_m_ of ATP in dose-response (9-pt curve in duplicate) for each kinase for which it was offered. Eurofins kinase enzymatic LANCE assay was carried out at the K_m_ of ATP for MAP2K5/MEK5 at a single concentration (10 μM) in duplicate.

### Cell Culture

HEK293 and HEK293T cells were acquired from the American Type Culture Collection (ATCC) and then cultured in a humidified incubator at 37°C and 5% CO_2_. Cells were passaged regularly using 0.05% Trypsin/0.53mM EDTA in Sodium Bicarbonate (Corning, 25-052-CI), and maintained in Dulbecco’s Modified Eagle Medium (DMEM) (Corning, 10-013-CV) supplemented with 10% fetal bovine serum (FBS). Analogous cell culture conditions have been previously employed by our groups^2,22^.

### NanoBRET measurements

NanoBRET assays were executed as described previously^22^. Constructs for BRSK2 NanoBRET measurements were provided in kind by Promega. The *N*-terminal Nanoluciferase (NL)/BRSK2 fusion (NL-BRSK2) was encoded in pFN32K expression vector, including flexible Gly-Ser-Ser-Gly linkers between NL and BRSK2 (Promega). For cellular BRSK2 NanoBRET target engagement experiments, a 10 μg/mL solution of DNA in Opti-MEM without serum (Gibco) was prepared with 9 μg/mL of Carrier DNA (Promega) and 1 μg/mL of NL-BRSK2 to achieve a total volume of 1.05 mL. To this solution was then added 31.5 μL of FuGENE HD (Promega) to form a lipid:DNA complex, mixed by inversion 8 times, and incubated at room temperature for 20 min. The resulting transfection complex (1.082 mL) was next gently mixed with HEK293 cells (21 mL) suspended at a density of 2 × 10^5^ cells/mL in DMEM containing 10% FBS. Finally, 100 μL of this solution was then dispensed into each well of a 96-well tissue culture treated plate (Corning, 3917), and the plate was incubated at 37°C with 5% CO_2_ for 24 hours.

After 24 hours, media was removed via aspiration and 85 μL of room temperature Opti-MEM without phenol red (Gibco) was added to each well. NanoBRET Tracer K5 (Promega) was used at a final concentration of 1 μM, the concentration previously determined to be optimal via a tracer titration experiment. Next, 5 μL (20x working stock of NanoBRET Tracer K5 [20 μM] in Tracer Dilution Buffer (Promega N291B)) was added to each well with the exception of the “no tracer” control wells. Test compounds were prepared as concentrated stock solutions in 100% DMSO (Sigma) at a concentration of 10 mM. They were then diluted in Opti-MEM media (99%) to prepare stock solutions containing 1% DMSO. A volume of 10 μL of 10-fold test compound stock solutions (final assay concentration of 0.1% DMSO) was added to each well. For “no compound” and “no tracer” control wells, 10 μL of Opti-MEM plus DMSO (9 μL Opti-MEM plus 1 μL DMSO) was added to each well to achieve a final concentration of 1% DMSO. The resultant 96-well plates containing transfected cells with NanoBRET Tracer K5 and test compounds (100 μL total volume per well) were equilibrated at 37°C with 5% CO_2_ for 2 hours.

After 2 hours, plates were returned to room temperature over the course of 15 min. To measure NanoBRET signal, a 3X stock solution was prepared by mixing NanoBRET NanoGlo substrate (Promega) at a ratio of 1:166 to Opti-MEM media in combination with extracellular NanoLuc Inhibitor (Promega) diluted 1:500 (10 μL [30 mM stock] per 5 mL Opti-MEM plus substrate). Next, 50 μL of the 3X substrate/extracellular NL inhibitor stock was added to each well. Finally, plates were read within 10 min of this addition using a GloMax Discover luminometer (Promega) equipped with 450 nm BP filter (donor) and 600 nm LP filter (acceptor), using 0.3 s integration time according to the “NanoBRET 618” protocol (Promega).

Test compounds were evaluated at 8 concentrations in competition with NanoBRET Tracer K5 in HEK293 cells transiently expressing the NL-BRSK2 fusion protein. Raw milliBRET (mBRET) values were obtained by dividing the acceptor emission values (600 nm) by the donor emission values (450 nm), and then multiplying by 1000. Averaged control values were used to represent complete inhibition (no tracer control: transfected cells in Opti-MEM + DMSO; tracer control: transfected cells in Opti-MEM + DMSO + Tracer K5 only), and were plotted alongside the raw mBRET values. The data with n=3 biological replicates was first normalized and then fit using Sigmoidal, 4PL binding curve in Prism Software (version 8, GraphPad, La Jolla, CA, USA).

### Plasmids and Reagents

ORFs used for Western blots were in a pHAGE-CMV-FLAG expression vector as previously reported^2^.

### Western blot

Western blots were carried out as described previously^2^. HEK293T cells were plated in 6-well format and incubated at 37°C with 5% CO_2_ for 12 hours. Next, cells were transfected with 1.5ug of plasmid per well for 24 hours and treated with GW296115 (in DMSO). Samples were lysed using standard conditions of RIPA (10% glycerol, 50mM Tris-HCL, 100mM NaCl, 2mM EDTA, 0.1% SDS, 1% Nonidet P-40, and 0.2% Sodium Deoxycholate) supplemented with protease inhibitor cocktail (ThermoFisher Scientific, 78429), phosphatase inhibitor cocktail (ThermoFisher Scientific, 78426), NEM (Thermo Scientific, 23030), and Benzonase (Sigma, E1014). Lysis was carried out on ice over the course of 30min and resultant lysates were centrifuged at 4°C for 15min at 21,000xg. Following normalization of protein concentration via BCA (Pierce, 23225), samples were denatured in NuPAGE LDS buffer (Invitrogen, NP0007) plus 1mM DTT. Blots were imaged with a LiCor Odyssey imager, and then quantified using ImageStudio 5.2. Finally, ANOVA with multiple comparison was performed for quantitated blots using Prism 8.2.4. All antibodies used are listed in **Table S3**.

## Data availability

All data generated or analyzed during this study are included in this published article (and its Supplementary Information File).

## Acknowledgements

The SGC is a registered charity (number 1097737) that receives funds from AbbVie, Bayer Pharma AG, Boehringer Ingelheim, Canada Foundation for Innovation, Eshelman Institute for Innovation, Genome Canada, Genentech, Innovative Medicines Initiative (EU/EFPIA) [ULTRA-DD grant no. 115766], Janssen, Merck KGaA Darmstadt Germany, MSD, Novartis Pharma AG, Ontario Ministry of Economic Development and Innovation, Pfizer, São Paulo Research Foundation-FAPESP, Takeda, and Wellcome [106169/ZZ14/Z]. Research reported in this publication was supported in part by the NC Biotech Center Institutional Support Grant 2018-IDG-1030, and by the NIH 1U24DK11604.

As part of the PKIS set, GW296115 was generously donated by GlaxoSmithKline to SGC-UNC. The Sorger lab carried out the deep dye drop assays. Constructs for NanoBRET measurements of BRSK2 were kindly provided by Promega. Coral was used to make the kinome trees depicted in Figures 1 and 6. Coral was developed in the Phanstiel Lab at UNC; http://phanstiel-lab.med.unc.edu/CORAL^23^.

## Contributions

C.W. and D.H.D. selected and sent GW296115 for *scan*MAX profiling at DiscoverX and deep dye drop analysis. A.D.A. selected kinases and sent GW296115 for follow-up at Eurofins. T.Y.T. and C.W. performed the cell-based studies. T.Y.T. and A.D.A wrote the paper with contributions from the rest of the authors. All authors have given approval to the final version of the manuscript.

## Additional Information

**Supplementary information** is available for this paper at (doi website)

**Correspondence** and requests for materials should be addressed to A.D.A.

## References

1 Elkins, J. M. et al. Comprehensive characterization of the Published Kinase Inhibitor Set. Nature biotechnology 34, 95–103, doi:10.1038/nbt.3374 (2016).

2 Tamir, T. Y. et al. Gain-of-function genetic screen of the kinome reveals BRSK2 as an inhibitor of the NRF2 transcription factor. Journal of Cell Science, jcs.241356, doi:10.1242/jcs.241356 (2020).

3 Uings, I. J., Spacey, G. D. & Bonser, R. W. Effects of the Indolocarbazole 3744W on the Tyrosine Kinase Activity of the Cytoplasmic Domain of the Platelet-Derived Growth Factor β-Receptor. Cellular signalling 11, 95–100, doi:https://doi.org/10.1016/S0898-6568(98)00039-4 (1999).

4 Wells, C. I. et al. The Kinase Chemogenomic Set (KCGS): An open science resource for kinase vulnerability identification. bioRxiv, 10.1101/2019.1112.1122.886523, doi:10.1101/2019.12.22.886523 (2019).

5 Davis, M. I. et al. Comprehensive analysis of kinase inhibitor selectivity. Nature biotechnology 29, 1046–1051, doi:10.1038/nbt.1990 (2011).

6 Metz, K. S. et al. Coral: Clear and Customizable Visualization of Human Kinome Data. Cell Syst 7, 347–350 e341, doi:10.1016/j.cels.2018.07.001 (2018).

7 Vasta, J. D. et al. Quantitative, Wide-Spectrum Kinase Profiling in Live Cells for Assessing the Effect of Cellular ATP on Target Engagement. Cell chemical biology 25, 206–214, doi:10.1016/j.chembiol.2017.10.010 (2018).

8 Lizcano, J. M. et al. LKB1 is a master kinase that activates 13 kinases of the AMPK subfamily, including MARK/PAR-1. EMBO J 23, 833–843, doi:10.1038/sj.emboj.7600110 (2004).

9 Bright, N. J., Thornton, C. & Carling, D. The regulation and function of mammalian AMPK-related kinases. Acta Physiol (Oxf) 196, 15–26, doi:10.1111/j.1748-1716.2009.01971.x (2009).

10 Wang, Y. L., Wang, J., Chen, X., Wang, Z. X. & Wu, J. W. Crystal structure of the kinase and UBA domains of SNRK reveals a distinct UBA binding mode in the AMPK family. Biochem Biophys Res Commun 495, 1–6, doi:10.1016/j.bbrc.2017.10.105 (2018).

11 Kim, J., Kundu, M., Viollet, B. & Guan, K. L. AMPK and mTOR regulate autophagy through direct phosphorylation of Ulk1. Nat Cell Biol 13, 132–141, doi:10.1038/ncb2152 (2011).

12 Daniel F. Egan, D. B. S., Maria M. Mihaylova,Sara Gelino, Rebecca A. Kohnz, William Mair, Debbie S. Vasquez, Aashish Joshi, Dana M. Gwinn, Rebecca Taylor, John M. Asara, James Fitzpatrick, Andrew Dillin, Benoit Viollet, Mondira Kundu, Malene Hansen, Reuben J. Shaw. Phosphorylation of ULK1 (hATG1) by AMP-Activated Protein Kinase Connects Energy Sensing to Mitophagy. Science (2011).

13 Wang, C. et al. Phosphorylation of ULK1 affects autophagosome fusion and links chaperone-mediated autophagy to macroautophagy. Nat Commun 9, 3492, doi:10.1038/s41467-018-05449-1 (2018).

14 Lim, J. et al. Proteotoxic stress induces phosphorylation of p62/SQSTM1 by ULK1 to regulate selective autophagic clearance of protein aggregates. PLoS Genet 11, e1004987, doi:10.1371/journal.pgen.1004987 (2015).

15 Ichimura, Y. et al. Phosphorylation of p62 Activates the Keap1-Nrf2 Pathway during Selective Autophagy. Molecular Cell 51, 618–631, doi:10.1016/j.molcel.2013.08.003 (2013).

16 Cloer, E. W. S., P. F.; Cousins, E. M.; Goldfarb, D.; Mowrey, D. D.; Harrison, J. S.; Weir, S. J.; Dokholyan, N. V.; Major, M. B. p62-Dependent Phase Separation of Patient-Derived KEAP1 Mutations and NRF2. Molecular and Cellular Biology, doi:DOI: 10.1128/MCB.00644-17 (2018).

17 Anastassiadis, T., Deacon, S. W., Devarajan, K., Ma, H. & Peterson, J. R. Comprehensive assay of kinase catalytic activity reveals features of kinase inhibitor selectivity. Nature biotechnology 29, 1039–1045, doi:10.1038/nbt.2017 (2011).

18 Bosc, N., Meyer, C. & Bonnet, P. The use of novel selectivity metrics in kinase research. BMC Bioinformatics 18, 17, doi:10.1186/s12859-016-1413-y (2017).

19 Zhao, B. et al. Structural basis for Chk1 inhibition by UCN-01. The Journal of biological chemistry 277, 46609–46615, doi:10.1074/jbc.M201233200 (2002).

20 Uitdehaag, J. C. M. et al. Comparison of the Cancer Gene Targeting and Biochemical Selectivities of All Targeted Kinase Inhibitors Approved for Clinical Use. PloS one 9, e92146, doi:10.1371/journal.pone.0092146 (2014).

21 G. Manning, D. B. W., R. Martinez, T. Hunter, S. Sudarsanam. The Protein Kinase Complement of the Human Genome. Science (2002).

22 Wells, C. et al. SGC-AAK1-1: A Chemical Probe Targeting AAK1 and BMP2K. ACS medicinal chemistry letters 11, 340–345, doi:10.1021/acsmedchemlett.9b00399 (2019).

23 Metz, K. S. et al. Coral: Clear and Customizable Visualization of Human Kinome Data. Cell systems 7, 347–350.e341, doi:10.1016/j.cels.2018.07.001 (2018).

