## Supplemental Information for "PKIS Deep Dive Yields a Chemical Starting Point for Dark Kinases and a Cell Active BRSK2 Inhibitor"

**Supplemental Table 1.** Select broad screening results around PKIS compound GW296115.

| Kinase | Nanosyn % Inhibition at 1 μM | TSA $\Delta T_m$ at 10 μM (°C) | IDG kinase |
|---|---|---|---|
| BRSK1 | 102 | NT | Y |
| BRSK2 | 95 | NT | Y |
| RSK3 | 92 | NT | N |
| STK17B/DRAK2 | NT | 18.3 | Y |
| PIM3 | 83 | 12.0 | N |
| PRKAA2 | 52[1] | 10.6 | N |
| STK33 | NT | 9.8 | Y |
| PIM2 | 68 | 9.8 | N |
| GAK | NT | 8.4 | N |

[1]Protein kinase AMP-activated catalytic subunit alpha 2

**Supplemental Table 2.** Select DiscoverX and Eurofins data generated for GW296115.

| Kinase | DiscoverX % Inhibition at 1 μM | >90%I at Nanosyn or >7.5°C $\Delta T_m$ in Table 1 | IDG kinase | Assay offered by Eurofins | Eurofins $IC_{50}$ (nM) | Family |
|---|---|---|---|---|---|---|
| STK17B/DRAK2 | 100 | Y (NS) | Y | Y | 5.5 | CAMK |
| LRRK2 | 100 | NT | N | Y | 25 | TKL |
| MAP3K19 | 100 | NT | N | N | NT | STE |
| PDGFRB | 99.8 | NT | N | Y | >10000 | TK |
| STK17A/DRAK1 | 99.6 | NT | Y | Y | 28 | CAMK |
| MAP2K5 | 99.5 | NT | N | Y | >10000 | STE |
| PLK4 | 99.5 | NT | N | Y | 17 | OTHER |
| PRKG2/PKG2 | 98.8 | NT | N | Y | 6.3 | AGC |
| TNIK | 97.7 | NT | N | Y | 10 | STE |
| DAPK3/ZIPK | 96.9 | NT | N | Y | 100 | CAMK |
| GAK | 96.7 | Y (TSA) | N | N | NT | OTHER |
| DAPK2 | 96.7 | NT | N | Y | 21 | CAMK |
| MINK1/MINK | 96 | NT | N | Y | 35 | STE |
| MYLK/MLCK/smMLCK | 95.4 | NT | N | Y | 49 | CAMK |
| MAP3K9/MLK1 | 95.1 | NT | N | Y | 26 | TKL |
| RPS6KA6/RSK4 | 94.7 | NT | N | Y | 12 | CAMK |
| PIM3 | 94.4 | Y (NS and TSA) | N | Y | 89 | CAMK |
| AURKC | 94.1 | NT | N | Y | 4100 | OTHER |
| CHEK2 | 92.5 | NT | N | Y | 2.6 | CAMK |
| MAP3K11/MLK3 | 92.4 | NT | N | Y | 1.5 | TKL |
| STK33 | 92 | Y (TSA) | Y | Y | 110 | CAMK |
| FLT3 | 91.5 | NT | N | Y | 37 | TK |
| PDGFRA | 91.4 | NT | N | Y | 710 | TK |
| MAP4K4/HGK | 91.4 | NT | N | Y | 29 | STE |
| SGK1/SGK | 91.2 | NT | N | Y | 710 | AGC |
| AURKB | 90 | NT | N | Y | 94 | OTHER |
| RIOK3 | 89 | NT | Y | N | NT | ATYPICAL |

| | | | | | | |
|---|---|---|---|---|---|---|
| SGK3 | 86 | NT | N | Y | >10000 | AGC |
| NUAK1/ARK5 | 85 | NT | N | Y | 15 | CAMK |
| EPHB6 | 85 | NT | N | N | NT | TK |
| RPS6KA3/RSK2 | 84 | Y (NS) | N | Y | 11 | CAMK |
| BRSK2 | 83 | Y (NS) | Y | Y | 8.4 | CAMK |
| DCLK3/DCAMKL3 | 83 | NT | Y | Y | 89 | CAMK |
| PIM2 | 83 | Y (NS and TSA) | N | Y | 10 | CAMK |
| BRSK1 | 82 | Y (NS) | Y | Y | 21 | CAMK |
| PHKG1 | 82 | NT | Y | Y | 20 | CAMK |
| MYLK3/caMLCK | 80 | NT | N | N | NT | CAMK |
| DAPK1 | 78 | NT | N | Y | >10000 | CAMK |
| ROS1 | 76 | NT | N | Y | >10000 | TK |
| AXL | 75 | NT | N | Y | 160 | TK |
| DMPK | 75 | NT | N | Y | >10000 | AGC |

NS: Nanosyn; NT: not tested; TSA: thermal shift assay

**Supplemental Table 3.** Antibodies.

| Antibody | Source | Reference # | Dilution (v:v) |
|---|---|---|---|
| **Mouse anti-FLAG** | Sigma | F3165 | 1:1000 |
| **Mouse anti-GAPDH** | Sigma | G8795 | 1:1000 |
| **Mouse anti-Vinculin** | Santa Cruz | sc25336 | 1:1000 |
| **Rabbit anti-phospho AMPK Substrate Motif [LXRXX(pS/pT)** | Cell Signaling Technologies | 5759 | 1:1000 |
| **Rabbit anti-AMPK** | Cell Signaling Technologies | 2532 | 1:1000 |
| **Rabbit anti-phospho AMPK T172** | Cell Signaling Technologies | 2535 | 1:1000 |
| **Rabbit anti-ULK1** | Cell Signaling Technologies | 4776 | 1:1000 |
| **Rabbit anti-phospho ULK1 S317** | Cell Signaling Technologies | 12753 | 1:1000 |
| **Rabbit anti-phospho ULK1 S757** | Cell Signaling Technologies | 6888 | 1:1000 |
| **Rabbit anti-SQSTM1/P62** | Bethyl | A302-856A | 1:1000 |
| **Rabbit anti-phospho SQSTM1/P62 S351** | MBL International | PM074 | 1:1000 |

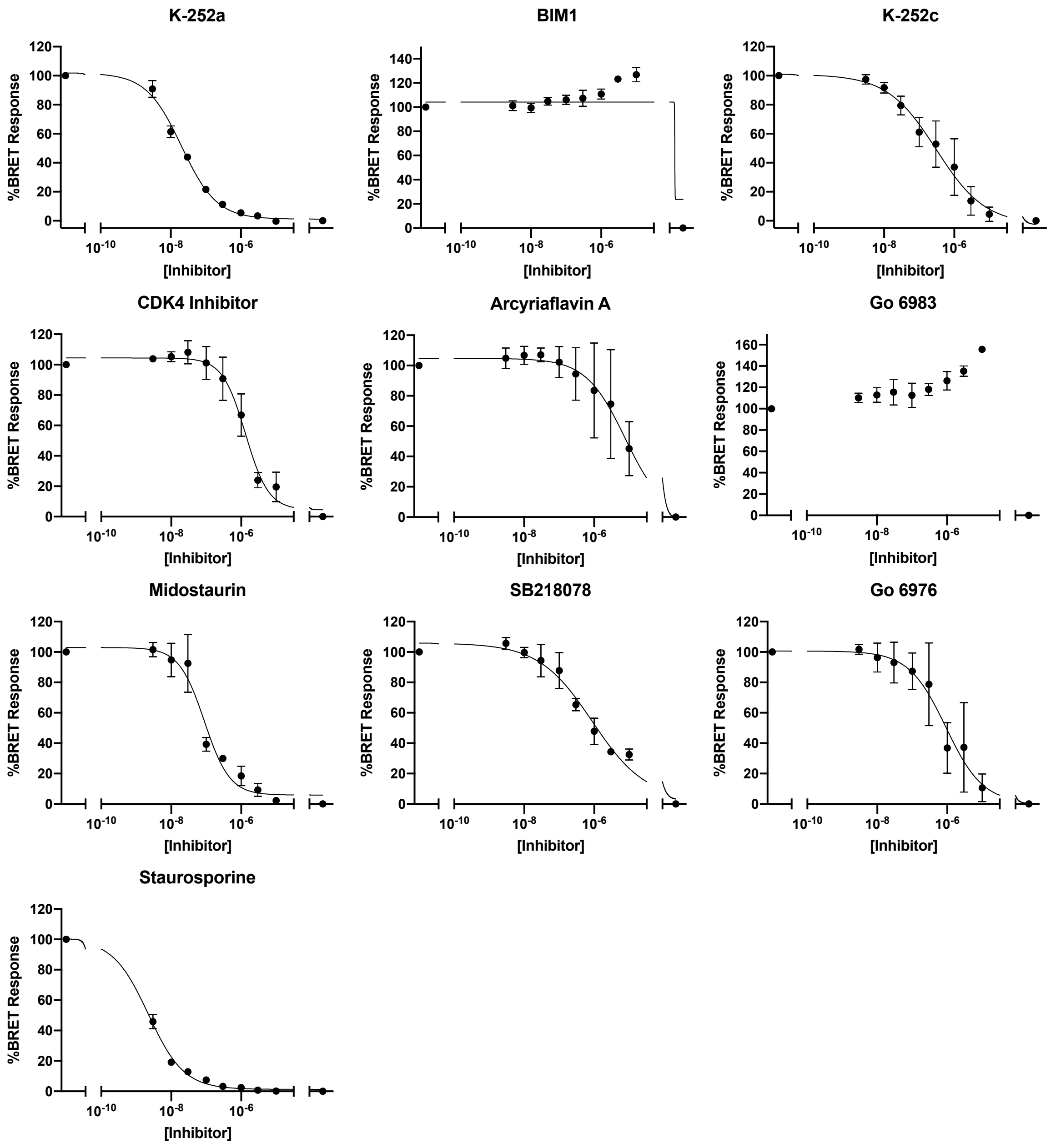


**Supplemental Figure 1.** NanoBRET cellular target engagement of indolocarbazoles and bisindolylmaleimides shown in **Fig. 5A** and **5B**.

### Figure 3B

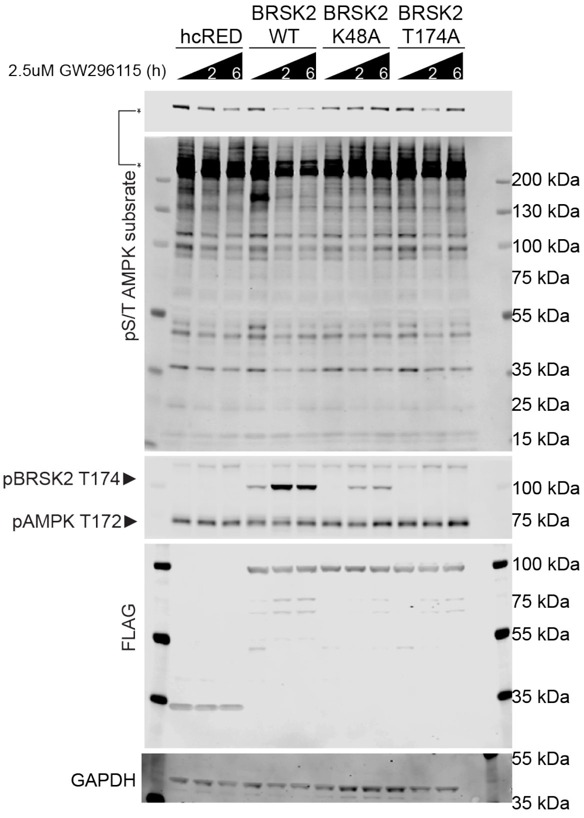


### Figure 4A

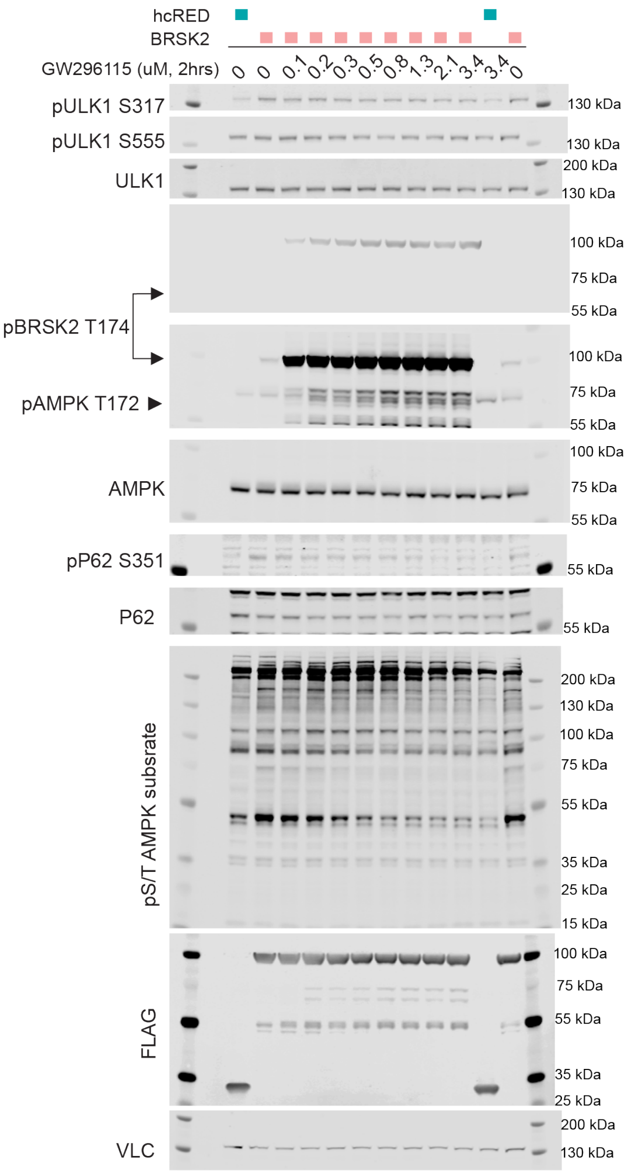


**Supplemental Figure 2.** Unformatted western blot images corresponding to **Fig. 3B** and **Fig. 4A**.
